# Medial Entorhinal VIP-expressing interneurons receive direct input from Anterior Dorsal Thalamus and are critical for spatial memory

**DOI:** 10.1101/2024.08.26.609578

**Authors:** Marie Oulé, Saishree Badrinarayanan, Rosa Sundar-Maccagno, Mark P. Brandon

**Affiliations:** Douglas Hospital Research Center, McGill University, Montreal, Canada; Integrated Program in Neuroscience, McGill University, Montreal, Canada; Department of Medicine, University of British Columbia

## Abstract

Head-direction (HD) cells are found across several regions in the brain, including the anterodorsal thalamic nucleus (ADN), the subicular complex, and the medial entorhinal cortex (MEC). A fundamental role of head direction cells is to provide input to MEC grid cells, which are thought to translate information about head direction into a metric code for spatial location. However, classic anatomical studies indicate that most thalamic HD projections pass indirectly to the MEC via the post- and para-subiculum, with only a small subset of ADN fibers terminating in the MEC. To further investigate the smaller and direct projection to the MEC, we use rabies-mediated retrograde tracing in mice to determine if this projection explicitly targets a subset of MEC neurons. Our findings reveal that ADN neurons specifically project onto MEC interneurons, with a preference for MEC VIP-expressing cells. Additionally, MEC VIP cells receive input from the hippocampus, the subicular complex, and the retrosplenial cortex - key centers for spatial memory - suggesting a specialized role for MEC VIP cells in spatial memory. Indeed, we find that MEC VIP cells exhibit increased c-Fos expression in a spatial memory task and show that chemogenetic inhibition of these neurons impairs task performance. Together, these data uncover a specific projection of head direction information onto MEC interneurons and confirm that MEC VIP-expressing cells are critical for spatial memory.

## Introduction

The medial entorhinal cortex (MEC) is essential for spatial navigation and memory^1–3^. Within the MEC microcircuit, excitatory cells in the superficial LII-LIII MEC layers exhibit various profiles of spatial firing^3^, such as grid cells^4^, boundary-vector cells^5,6^, head-direction cells^6,7,8^, or object-vector cells^9^, whose activity support the neural representation of space. Among these spatially-tuned cells, head-direction cells provide critical inputs to cortical grid cells ^10,11^, and the activity of these two cell types is essential for the emergence of the cognitive map of an environment and necessary for spatial memory. Head direction tuning is present across various cortical and subcortical limbic regions such as the anterodorsal nucleus (ADN) of the thalamus^12,13^, the retrosplenial cortex^14^, the subicular complex^15,16^, and the entorhinal cortex^17,18^. Although these regions are highly interconnected^17^, the propagation of the head direction information is thought to follow a preferential direction^19^ in which the thalamic head direction signal propagates to the MEC through the para- and pre-subiculum^1,20,21^. Indeed, former neuroanatomical studies have shown that most ADN axons target the subicular complex, but few projections have been reported in the MEC^17,22^. It remains possible that the sparse, direct projections from ADN to the MEC carry critical information targeted to specific MEC sub-populations.

While tremendous advances have been made in understanding excitatory cells at the physiological^23–26^, functional^23–27^, and behavioral^23–25^ levels, less is known about how interneurons regulate the information flow necessary for the emergence of space coding in the MEC^23–25,28–30^. The MEC is rich in inhibitory cells, such as parvalbumin (PV)^28,31^, somatostatin (SOM)^31,32^, or Vasoactive Intestinal Peptide (VIP) cells^33^, embedding the excitatory cells^28,31,34^ in a network of inhibition. In contrast to PV and SOM cells, which directly project onto excitatory cells^1,30^, VIP cells are mostly disinhibitory^35–38^. These interneurons either project onto PV or SOM cells and thus indirectly orchestrate excitatory cells’ activity by releasing them from ongoing inhibition. In the MEC, pharmacogenetic inactivation of PV cells resulted in impairment in the periodicity of grid cells^28^. In contrast, the inactivation of SOM cells altered the spatial selectivity of aperiodic cells^28^ but left border or head-direction activity unaffected^28^. While MEC VIP cells’ firing and morphological characteristics have been reported^33^, less is known about their role in entorhinal spatial coding properties. Despite limited knowledge, recent work highlights VIP cells as critical for memory in the auditory cortex^37^, the amygdala^35^, and the hippocampus^39,40^. Interestingly, recent evidence demonstrates that hippocampal VIP cells are functionally versatile as they do not only modulate principal cells’ firing but adapt their activity according to task demands^39^, supporting the hypothesis that interneurons integrate external information for spatially-guided behavior. Moreover, VIP cells are activated by diverse long-range inputs in the prefrontal cortex ^41^, the anterior insular cortex^42^, and the amygdala^35^. Hence, we hypothesized that MEC VIP cells, as observed in other brain regions, would receive external inputs from brain regions involved in space processing, such as the ADN, and would therefore be necessary for cognitive function supported by the MEC, such as spatial memory.

## Results

### VIP cells receive inputs from key memory and space brain centers

VIP^cre^ mice were first unilaterally injected with the helper virus AAV2/8-hSyn-FLEX-TVA-P2A-eGFP-2A-oG (TVA-eGFP) in the MEC and four weeks later received the RABV-DeltaG-EnvA-mCherry rabies virus (Fig. 1A). This procedure allowed for the localization and quantification of input cells (mCherry+), which made synaptic connections with the starter cells (GFP+/mCherry+) (Fig. 1B). The viral approach used showed a high level of expression (Fig. 1G) and specificity (Fig. 1H) as the TVA-eGFP virus was expressed in all VIP/TdTom cells near the injection sites, and no leakage was observed in VIPcre-negative animals injected with the TVA-eGFP virus (Fig. 1H). For quantification analysis, we included brain regions with at least one input cell in both planes (identified by the presence of mCherry+ cell bodies). The relative quantification of inputs received by individual starter cells (called convergence index (CI)) was calculated by dividing the number of MEC starter cells by the number of input cells in a given brain region. VIP starter cells were observed across all the layers of the MEC, with a majority of them localized in the superficial layers and less in the deep layers (Fig. 1C, suppl. Table 1, superficial: 48.75 ± 10.94, deep: 17 ± 5.672). This observation is consistent with a previous report identifying more VIP cells in the superficial layers^33^. Analysis showed that VIP cells are highly connected within MEC microcircuitry as most of the input cells were located within the structure (Fig. 1D, suppl. Table 1, CI = 21.56 ± 9.049,), especially the superficial layers (superficial CI = 15.28 ± 6.382, deep CI = 6.279 ± 2.692). These cells also received large projections from CA1 (Fig. 1E, F, I and J, suppl. Table 1 CI = 8.702 ± 4.886) and the subicular complex (Fig. 1E, F, I and I, suppl. Table 1 CI = 5.486 ± 1.551). Among the cortical regions, dense afferents originating from the perirhinal (Fig. E and K CI = 0.3947 ± 0.3828) and the retrosplenial (Fig. 1F, J and K, CI = 1.685 ± 0.8703, 5.1% of the total input cells) cortices were found. In contrast, sparse afferents came from the lateral entorhinal cortex (LEC) (Suppl. Fig. 3). Overall, projections from cortical areas known to process spatial memory constituted the largest population of projecting neurons (Fig. 1H, MEC: 41.6%, CA1 25.5%, and subicular complex 11,3%) as 78.4% of labeled soma were located in one of these regions. In the forebrain, input cells were also identified in the medial septum/horizontal diagonal band (Fig. 1K and L, suppl. Table 1, CI = 0.2586 ± 0.1284). While sporadic input cells were found in the thalamus, a strikingly significant population of input cells was located in the ADN (Fig. 1J and 2E, CI = 1.934 ± 0.7904). Nevertheless, ADN contributes to 4.7% of the total projections received by MEC VIP cells (Fig. 1J). Residual input cells were also observed from the amygdala or the visual and auditory cortices (Fig. 1K). This result demonstrates that MEC VIP cells are highly interconnected within the MEC microcircuit. Meanwhile, this cell population receives a significantly wider spectrum of inputs, including major head-direction coding centers such as the ADN and the subicular complex, but also regions essential to memory such as the hippocampus, the retrosplenial, and the perirhinal cortices.

**Figure 1.**
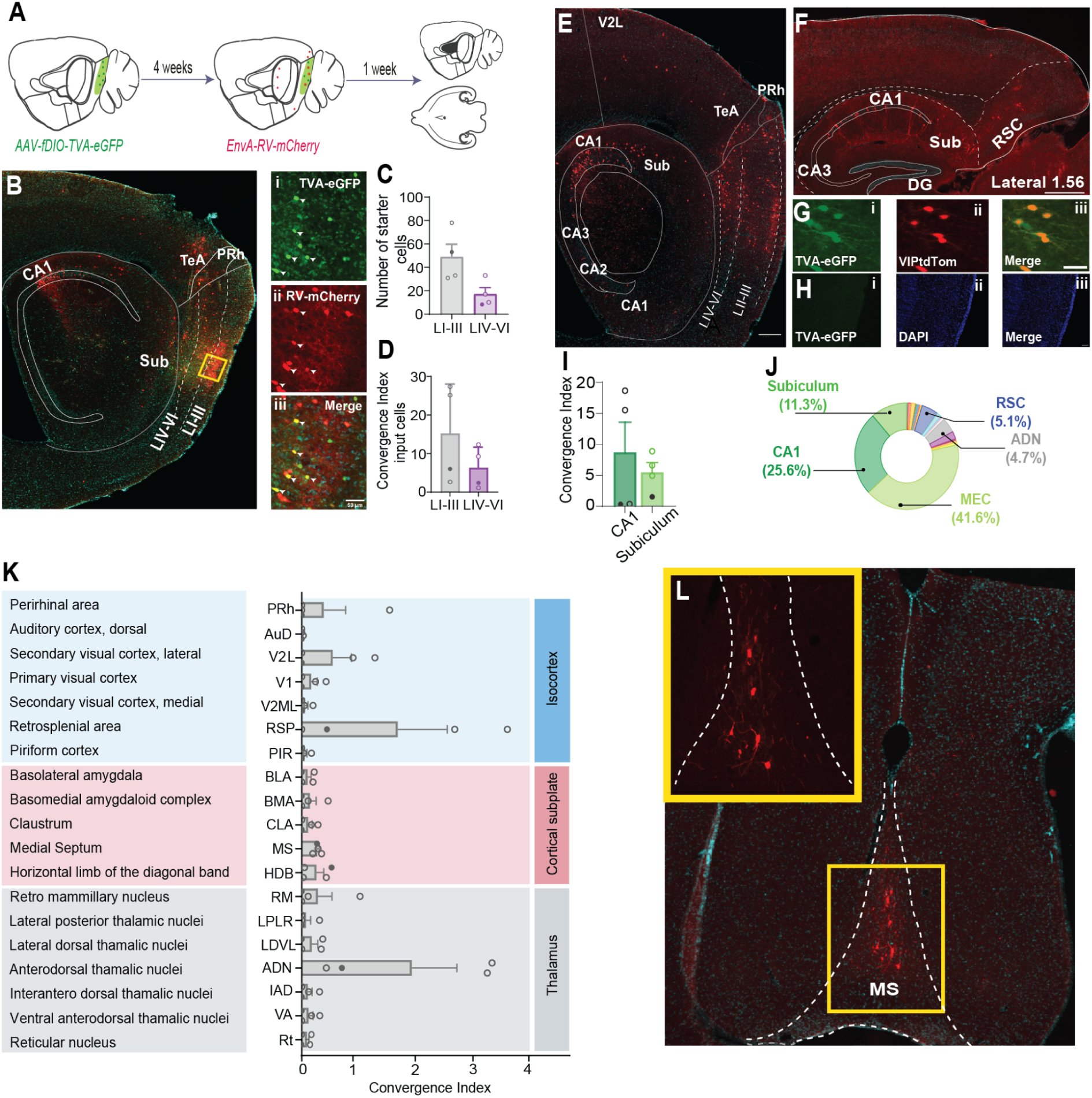
Tracing local and long-range projections to VIP cells in the MEC A. Strategy for helper virus and rabies virus injection to enable retrograde tracing of inputs to the MEC. The helper virus (TVA-eGFP) was injected into a VIP-cre mouse first, and then four weeks later, the rabies virus (RV-mCherry) was injected. The animals were perfused a week later for whole brain sectioning. B. Image of the injection site and starter cells in VIP^cre^ mouse. Dashed lines mark layers of the MEC. Yellow boxes and white arrows denote a magnified view of the starter and input cells within the MEC. Transfection of VIP cell by (i) TVA receptors (TVA-eGFP) and (ii) EnvA protein (mCherry). Colocalization of the two channels is seen in (iii) the starter cell. C. Quantification of VIP starter cells in the MEC represented as the number of starter cells across the different layers of the MEC D. Quantification of input cells within different layers of the MEC relative to the total number of starter cells. E. Representative image of monosynaptic input cells from different layers of the MEC, secondary visual cortex, perirhinal cortex, and the hippocampal formation F. Representative image of inputs arising from the CA1 region of the hippocampus, subicular complex, and retrosplenial cortex. G. Representative images of the TVA-eGFP virus injected in VIPTdTom mice to confirm that the TVA-eGFP virus is well expressed in VIP interneurons. H. Representative images of the TVA-eGFP virus injected in the VIP^cre^-negative animal show no viral expression leakage. I. Input cell quantification for cells projecting from the CA1 and subicular complex J. The total fraction of inputs from different brains projecting to VIP cells in the MEC K. Input cell quantification for different brain regions in the isocortex, cortical subplate, and thalamus, as defined by the Allen Brain Atlas. The closed circle indicates sections that were sliced in the horizontal plane L. Representative images of inputs arising from the medial septum (MS) The CI for all graphs was calculated as a ratio of the number of inputs from a given brain region over the number of starter cells quantified in an animal. VIP^cre^ animals n = 4. In all panels C, D, I and K the closed circle indicates sections that were sliced in the horizontal plane

**Figure 2.**
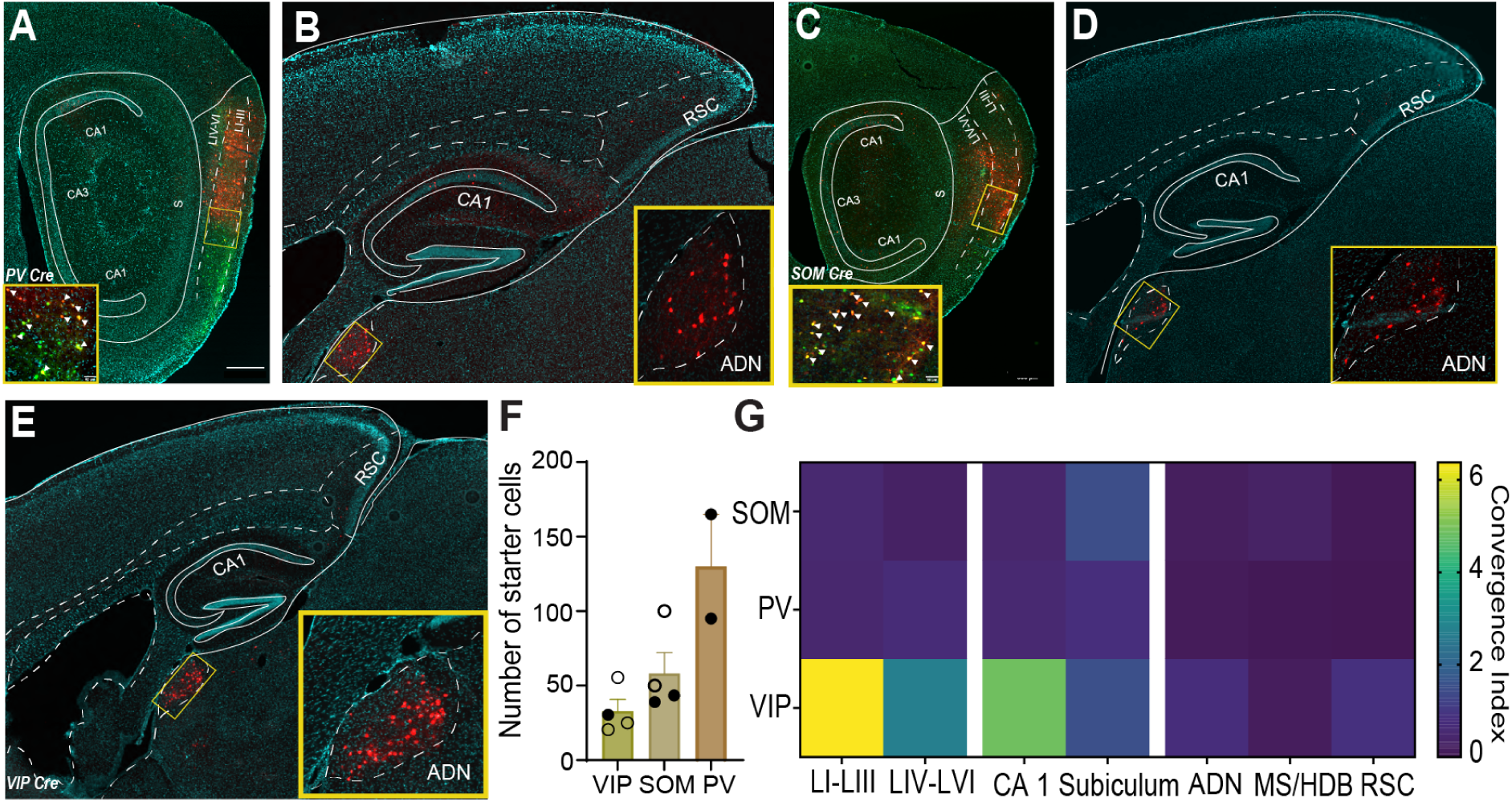
Interneurons in the MEC receive projections from the ADN A. Representative image of starter cells in PV^cre^ animals. Starter cells are highlighted in yellow and indicated by white arrowheads. Scale bar in the inset is 100 microns. B. Representative image displaying input cells in the ADN and RSC of PV^cre^ mice injected with rabies-mediated retrograde virus. C. Representative image of starter cells in SOM^cre^ animals. Starter cells are highlighted in yellow and indicated by white arrowheads. The Inset scale bar represents 100 microns. D. Representative image displaying input cells in the ADN SOM^cre^ mice injected with the rabies-mediated retrograde virus. E. Representative image displaying input cells in the ADN VIP^cre^ mice injected with the rabies-mediated retrograde virus. F. Quantification of starter cells in the MEC for different interneuron populations. The closed circle indicates sections that were sliced in the horizontal plane G. Heat maps illustrate the quantification of the inputs to VIP, PV, and SOM MEC interneurons based on the convergence index for each cell line: VIP^cre^ (n = 4 mice), SOM^cre^ (n = 4 mice), and PV^cre^ (n = 2 mice). ADN - Anterodorsal thalamic nucleus, MS - Medial Septum, HDB - Horizontal Diagonal Band of Broca, and RSC - Retrosplenial cortex. VIP^cre^ animals n = 4, SOM^cre^ n = 4, PV^cre^ n = 2. In all panels the closed circle indicates sections that were sliced in the horizontal plane

### Thalamic ADN projections target MEC interneurons, with a preference for VIP cells

Direct ADN-MEC connections were reported to be very sparse ^22^ compared to projections to adjacent structures^20,21^, leading to the consensus that head direction information predominantly arrives at the MEC via the pre- and para-subiculum. To determine the extent to which this connectivity pattern is specific to VIP cells, the same experimental approach was applied to localize thalamic neurons projecting onto other MEC interneurons, such as PV (Fig. 2A, B and suppl. Fig. 1B) and SOM (Fig. 2C, D and suppl. Fig. 1C), as well as stellate (Fig. 3A, B and suppl. Fig. 2) and superficial pyramidal cells (Fig. 3C, D and suppl. Fig. 2). The greatest number of starter cells was observed in the PV^cre^ line (Fig. 2F, 122.5 ± 42.5), followed by SOM^cre^ (Fig. 2F, 58.13 ± 14.14) and VIP^cre^ lines (Fig. 2F, 32.88 ± 7.814). This distribution is consistent with other studies showing that PV is the largest medial entorhinal cortical interneuron subtype^43,44^. Strikingly, ADN’s projection showed a strong preference towards VIP over SOM and PV cells (Fig. 2G, suppl. Table 1 CI = 1.934 ± 0.709, CI = 0.2667 ± 0.1568 and CI = 0.2915 ± 0.0415 respectively). Moreover, ADN cells connecting onto entorhinal cortical pyramidal and stellate cells were negligible or absent (Fig. 3F and suppl. Fig. 2 and Table 1, pyramidal cells: starter cells = 88.13 ± 51.15, CI = 0.02008 ± 0.008367; stellate cells: starter cells = 62 ± 25.66, CI = null). The sparsity of ADN projections onto MEC might result from their selectivity for entorhinal inhibitory cells. Although VIP cells represent the smallest interneuron population in the MEC, this result suggests that this cell population might be the principal direct relay of anterodorsal thalamic head-direction signal to the MEC.

**Figure 3:**
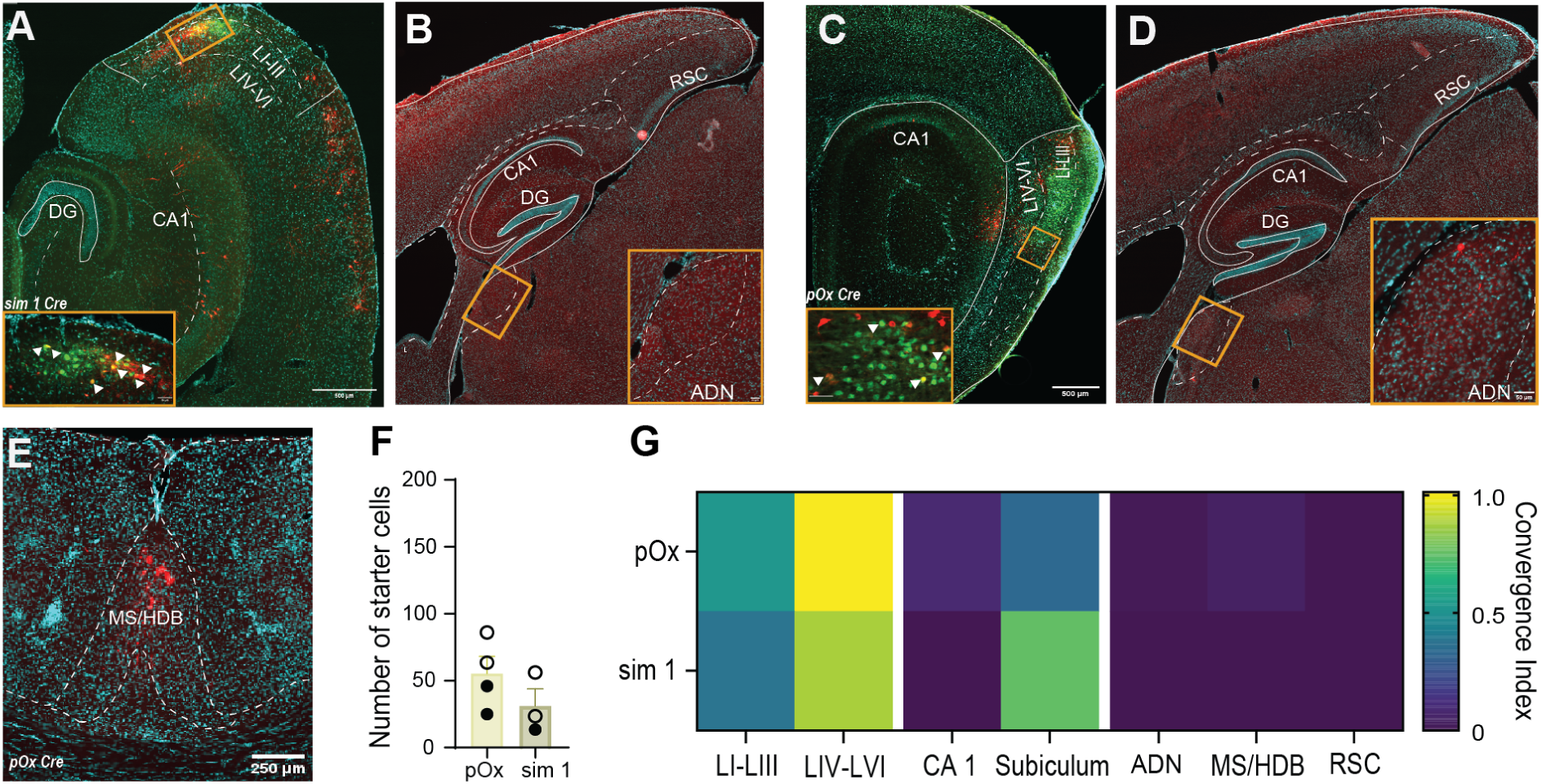
Tracing of inputs to excitatory cells in the MEC A. Representative image of starter cells in sim1^cre^ animals. Starter cells are highlighted in yellow and indicated by white arrowheads. B. Representative image of the absence of input cells in the ADN and RSC of sim1^cre^ mice injected with rabies-mediated retrograde virus. C. Representative images of starter cells in pOx^cre^ animals. Starter cells are highlighted in yellow and indicated by white arrowheads. D. Representative image of the absence of input cells in the RSC of pOx^cre^ mice injected with rabies-mediated retrograde virus. Few input cells could be identified in the ADN of pOx^cre^ mice. ADN - Anterodorsal thalamic nucleus, MS - Medial Septum, HDB - Horizontal Diagonal Band of Broca, and RSC - Retrosplenial cortex. E. Representative image of input cells in the MS/HDB region in pOx^cre^ animals. F. Quantification of starter cells in the MEC for different excitatory cell populations. Full circles indicate sections sliced in the horizontal plane, and empty circles indicate sections sliced in the sagittal plane. G. Heat maps depict input tracing to sim1^cre^ and pOx^cre^ MEC excitatory cells. Each row displays the convergence index for each cell line: sim1^cre^ (n = 3 mice) and pOx^cre^ (n = 4 mice).

### VIP cells receive more intra-and inter-regional projections than the other medial entorhinal cellular subtypes

Within multiple subtypes in MEC, VIP cells appeared to receive the densest projections from other entorhinal cortical cells (Fig. 2G and suppl. Fig.1A and Table 1, VIP CI = 21.56 ± 9.049, PV CI = 3.62 ± 0.8701; SOM CI = 3.404 ± 0.6018; Fig. 3F and suppl. Fig. 2, pyramidal cells CI = 2.335 ± 0.7873; stellate cells CI = 4.051 ± 0.9715). We observed similar patterns for projections from CA1 (Fig. 2G and suppl. Fig. 2, VIP CI = 8.702 ± 4.886; PV CI = 1.684 ± 0.4963; SOM CI = 1.652 ± 0.4143; Fig. 3F and suppl. Fig. 2, pyramidal cells CI = 0.334 ± 0.08654; SC: null), the subicular complex (Fig. 2G and suppl. Fig. 2, VIP CI = 5.486 ± 1.551; PV CI = 2.76 ± 0.39; SOM CI = 2.266 ± 1.48; Fig. 3F and suppl. Fig. 2, pyramidal cells CI = 1.028 ± 0.3369; stellate cells CI = 1.815 ± 0.751), and the retrosplenial cortex (Fig. 2G and suppl. Fig. 2, VIP CI = 1.685 ± 0.8703; SOM CI = 0.03831 ± 0.03831; PV: null, Fig. 3F and suppl. Fig. 2, pyramidal and stellate cells: null), from which inputs were only observed onto VIP and SOM cells. Inputs from the medial septum/horizontal diagonal band made contact with all cell types, except stellate cells (Fig. 2G, Fig. 3E and G and suppl. Fig. 2), with a slight preference for SOM cells (Fig. 2G and suppl. Fig. 2 VIP CI = 0.2586 ± 0.1284; PV CI = 0.07424 ± 0.02576; SOM CI = 0.3306 ± 0.2934; Fig. 3F and suppl. Fig. 1 pyramidal cells CI = 0.007252 ± 0.00475; stellate cells: null). Perirhinal cortex inputs to MEC cells were also very sparse and restricted to VIP, SOM, and stellate cells, the latter receiving the most inputs (Fig. 2G and suppl. Fig. 1, VIP CI = 0.3947 ± 0.3828, SOM CI = 0.04981 ± 0.04981, PV: null, Fig. 3F and suppl. Fig. 1, stellate cellsCI = 0.5114 ± 0.1923, pyramidal cells: null). Regarding the LEC, no input cells were found in any interneuron mice lines, but some inputs projecting onto pyramidal and stellate cells were spotted (suppl. Fig. 3).

### VIP cells are activated by spatial changes in the environment and are essential for spatial memory

To assess the putative role of VIP cells in spatial memory, VIPtdTom mice were assayed on the MEC-dependent novel object location (NOL) task (Fig. 4A)^45,46,23,47^. As previously reported, mice could discriminate an object’s new location in the dissimilar condition but not in the extra-similar condition (Fig. 4B, dissimilar 0.3328 ± 0.04026 p<0.0001; extra-similar: −0.0215 ± 0.05194, p = 0.6933; dissimilar vs. extra-similar: p = 0.0002). Differences in discrimination ratio were not driven by exploration time during the sampling (Fig. 4C) or sex (suppl. Fig. 4A). To determine whether VIP cells were activated during the NOL task, we imaged the c-Fos immediate early gene^48^ 90 min after the test phase (Fig. 4A). No difference in the c-Fos population was found between control and test conditions when looking at the total number of c-Fos+ cell (Fig. 4E, behavioral control: 13132 ± 2041; dissimilar: 17214 ± 1813; extra-similar: 18214 ± 1490; p = 0.1453) or superficial vs. deep layers (Fig. 4E, superficial: behavior control 10374 ± 1841, dissimilar 13741 ± 1622, extra-similar 15290 ± 1432, p = 0.1355; deep: behavior control 2923 ± 433, dissimilar 3590 ± 633, extra-similar = 3169 ± 626.3, p = 0.6973). However, the number of VIP cells expressing c-Fos (VIP/c-Fos+cells) increased when animals were engaged in a discrimination-based task (Fig. 4F, behavior control 13 ± 1.169, dissimilar 24.49 ± 3.234, extra-similar 27.65 ± 1.789; p = 0.0006, behavior control vs. extra-similar: p = 0.0008, behavior control vs. dissimilar: p = 0.0052), independent of the behavioral conditions (extra-similar vs. dissimilar: p = 0.58). Moreover, the effect of behavior on the number of VIP/c-Fos+ cells was specific to superficial layers (Fig. 4F, behavior control 12.44 ± 1.499, dissimilar 25.1 ± 5.049, extra-similar 29.72 ± 3.677; p = 0.0093, behavior control vs. extra-similar: p = 0.0095; behavior control vs. dissimilar: p = 0.0549, dissimilar vs extra-similar: p = 0.6465), as no differences were observed when analyzing the deep layers only (Fig. 4F, behavior control 14.83 ± 6.693, dissimilar 40.05 ± 13.6, extra-similar 19.33 ± 7.476; p = 0.1742). Altogether, these results demonstrate that spatial memory is associated with the activation of VIP cells located in the superficial layer, independent of behavioral performance.

**Figure 4:**
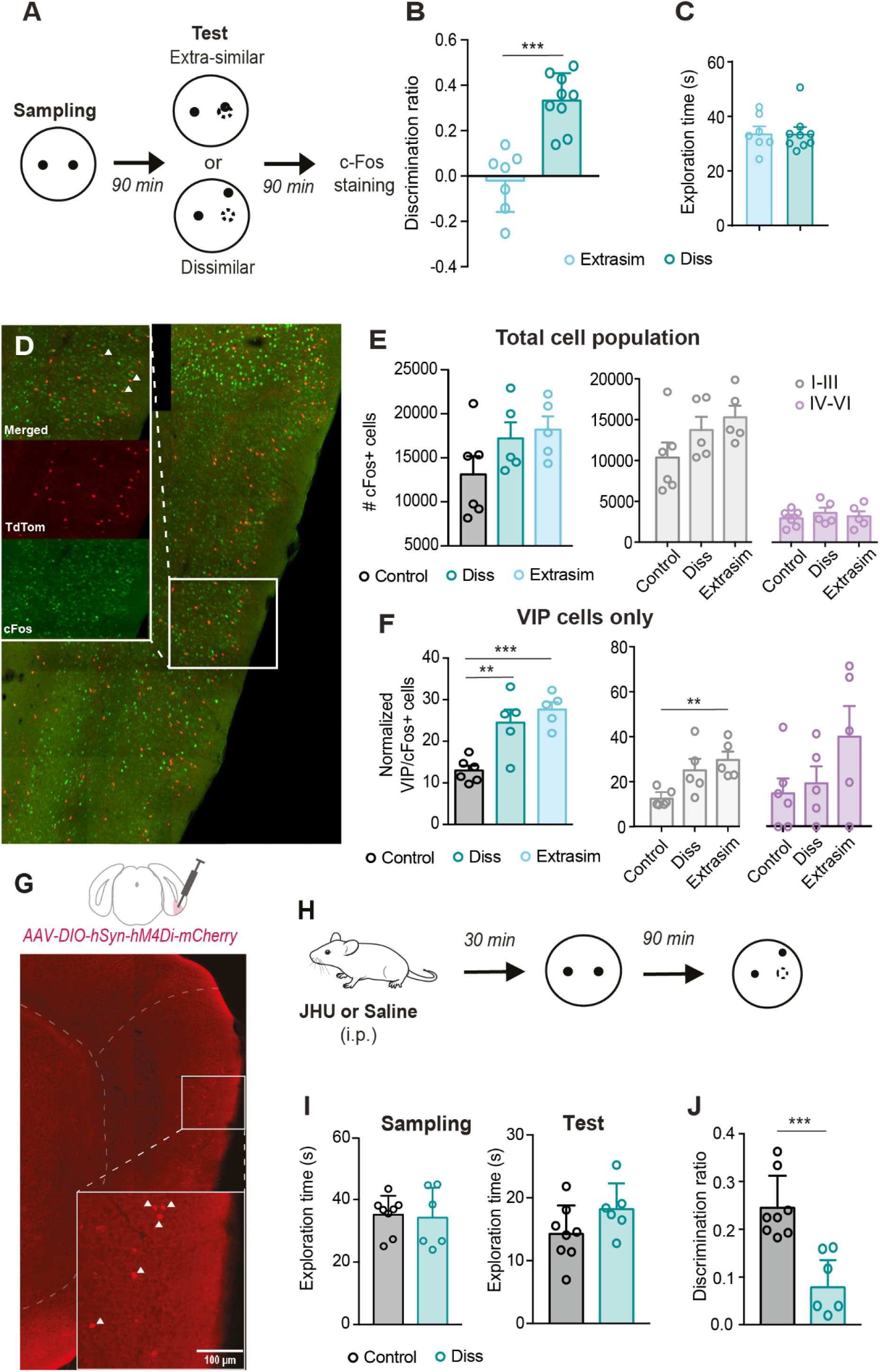
VIP interneuron recruitment is sensitive to changes in spatial environment and is necessary for spatial memory. A. Schematic of the novel object location (NOL) task. B. VIP-cre mice can significantly discriminate a new object’s location in the dissimilar, but not the extra-similar, version of the NOL task (extrasimilar n = 6, dissimilar n = 9, *** p<0.001). C. VIP-cre mice in the extra-similar and dissimilar conditions spend similar time exploring objects during the sampling phase. D. c-Fos immunostaining illustration on MEC section from VIP-TdTom mice. E. Quantification of the c-Fos+ cells population across all MEC layers (left panel) or in superficial vs. deep MEC layers (right panel). Behavior control n = 6, dissimilar n = 5, extra similar n = 5. F. Quantification of the VIP/c-Fos+ cells across all the MEC layers (left panel) or in superficial vs. deep MEC layers (right panel) (**p<0.01; ***p<0.001). G. Inhibitory AAV-DIO-hSyn-hM4Di-mCherry virus was injected unilaterally into the MEC of VIP^cre^ mice. H. JHU (1mg/kg) or saline was administered 30 min before the sampling phase. I. No difference in the exploration time during the sampling (left panel) and test (right panel) was observed between the control (n = 7) and JHU-administered group (n=6). J. Mice that received JHU prior sampling show a significant impairment in their ability to discriminate an object’s new location (***p<0.001). All data are represented with ± SEM.

To elucidate the role of VIP cells in spatial memory, VIP^cre^ mice received unilateral injections of AAV8.hSyn.DIO.hM4Di.mCherry virus and later were injected with JHU^49^ to drive the selective inhibition of VIP cells (Fig. 4G). Two control groups were used: the JHU control group did not receive viral injection but did receive JHU before behavior, while the virus control group did receive an injection of the AAV8.hSyn.DIO.hM4Di.mCherry virus but received saline instead of JHU, before sampling. Although a slight but significant increase in the exploration time was observed in the virus control group compared to the JHU control group (suppl. Fig. 4B, p = 0.0497), no difference in the behavioral performance was detected between groups (suppl. Fig. 4C, p = 0.4803). Therefore, both groups were pooled together and referred to as the behavioral control group (Fig 4I, J). As JHU was injected 30 minutes before the sampling, VIP cell activity was reduced throughout the behavioral procedure. Although JHU injection did not entirely abolish the ability of mice to detect an object’s new location in the dissimilar condition (Fig. 4J, behavior control: 0.2451 ± 0.02369, p < 0.0001; dissimilar: 0.07882 ± 0.02322, p = 0.0194), JHU severely altered the ability of mice to perform the task (Fig. 4J, p = 0.0004). This effect could not be explained by changes in the exploratory behavior as the time spent exploring objects was similar between groups for both the sampling (Fig. 4I, behavior control: 35167 ± 2137, JHU: 34210 ± 3900, p = 0.8220) and the test (behavior control: 14266 ± 1591, JHU 18201 ± 1668, p = 0.1184). Similar results were observed when mice received bilateral virus injection (suppl. Fig. 4D and E, exploration time sampling, unilateral: 34210 ± 3900, bilateral: 38099 ± 8039, p = 0.7619; discrimination ratio, unilateral: 0.07882 ± 0.02322, bilateral: 0.006875 ± 0.0893, p = 0.7619). Hence, this result demonstrates that the inactivation of MEC VIP cells impairs discrimination between similar spatial memories.

## Discussion

Most of our understanding of the biological mechanisms supporting MEC’s ability to create spatial representations comes from the study of principal cells. However, the role of interneurons in shaping spatial representations found in the MEC remains unclear. First, our study corroborates similar results obtained in various brain regions supporting the idea that VIP cells are important to relay information provided by long-range afferents into the local network, as observed in other brain regions such as the amygdala ^35^ or in the anterior insular cortex. More specifically, this report sheds new light on the critical role of VIP cells in MEC connectivity to distributed regions known to play central roles in spatial representation, and memory function. First, we demonstrate that MEC interneurons, particularly those that are VIP-expressing, are densely connected not just to other MEC cells but also to the hippocampus, the medial septum, the subicular complex, and the retrosplenial cortex. Most surprisingly, retrograde tracing analysis unveiled the existence of a direct thalamocortical pathway in which the ADN sends direct projections specifically to the MEC inhibitory network. Importantly, VIP cells are more densely connected to thalamic inputs, suggesting that these cells play a key role in relaying a thalamic head-direction code to other MEC cells. We then show that VIP cells were activated when spatial changes were made to the environment. Finally, we found that chemogenetic inhibition of VIP cells profoundly impaired the ability of mice to discriminate between similar memories.

### ADN projections to the MEC are selective to the inhibitory network and essentially target VIP cells

This study reports a unique selective projection from the ADN to MEC interneurons, where most anterodorsal thalamic inputs target VIP cells. Such an observation challenges a popular view on the propagation of the head-direction signal from the thalamus to cortical regions, wherein the MEC is thought to indirectly receive thalamic HD signal through superficial layers of the presubiculum^15,16,50^. Moreover, MEC-projecting pyramidal cells in the presubiculum receive direct projections from the ADN^15,16,50^, suggesting a disynaptic connection between the ADN and the MEC. As our study reports a direct ADN-MEC pathway, this raises several questions: 1) Do VIP cells receiving thalamic HD signal modulate the same population of cortical HD cells as the presubiculum or a different subpopulation? 2) Are these two pathways activated in the same context, or do they depend on environmental cues and features? 3) How do these two pathways modulate HD computation in the MEC? Future work should aim to identify cell populations and pathways supporting HD signal propagation and modulation in the MEC.

The role of an inhibitory drive in the head-direction signal is relatively limited. Investigations on this topic are mostly restricted to studies in the presubiculum^15,16,50^, where inhibitory cells control the information flow and shape the neuronal activity underlying HD coding. In this regard, it is interesting that neither inhibiting MEC PV nor SOM cells affected cortical HD cells^28^. In the MEC, approximately two-thirds of the VIP cells co-express calretinin^33^, a common marker for disinhibitory interneurons^51^. One hypothesis is that the remaining third of MEC VIP cells that do not express calretinin directly target excitatory cells in the MEC. The ADN-mediated activation of this population of VIP cells might participate in the shaping of HD activity in the MEC. However, in light of current evidence, we cannot rule out the possibility that some of the ADN inputs might also target excitatory cells in the MEC deep layers.

In addition, the number of thalamic afferents reaching a cell population isn’t the sole determinant of a pathway’s importance for target populations. Although ADN innervation to PV and SOM interneurons is sparse compared to the VIP cell population, the weight of this connection can be balanced by cellular and molecular plasticity properties facilitating or depressing the integration of thalamic information. To address this question, future work to record interneurons in physiological conditions during manipulation of ADN-MEC projections would be necessary to determine how the thalamic head-direction signal influences entorhinal cortical interneurons.

### VIP cells in memory: a ubiquitous mechanism for memory formation

The role of VIP cells in memory has been linked to their ability to release principal cells from ongoing inhibition, which is important for associative memory ^35,37^. Our findings on the effect of silencing MEC VIP cells in a spatial discrimination task further validate the significance of this cell group in memory processes. However, results obtained from c-Fos imaging suggest that VIP cells may have a more dynamic function, in addition to their role in associative memory. Specifically, the recruitment of VIP cells increased when changes were made to the spatial layout of objects, independent of any corresponding changes in behavioral response, suggesting a role of VIP cells in sensing changes in the environment. This observation resonates with recent works investigating discrimination mechanisms in the hippocampus. In a virtual environment where mice learn the location of a reward, small changes in the environment did not change the behavioral response of the animal, although DG’s cell activity changed upon exposure to a new, yet very similar, environment^52^. Interestingly, when the new environment was made more dissimilar, mice could learn the new location of the reward, with DG and CA1 areas showing altered activity in response to the dissimilar environment^52^. The result suggests that separate representations of the two environments were implemented throughout the hippocampus. This work demonstrates that (1) neural discrimination does not parallel behavioral discrimination and (2) that novelty leads to an adaptation in the behavioral response only if environmental changes are salient enough to modulate the activity of specific brain regions, such as CA1^52^, to allow the formation of representation of new experiences. A similar observation has been made in the prefrontal cortex, where ventral hippocampal inputs to prefrontal VIP-expressing cells drive different representations of the open and closed arms of an elevated-plus maze^41^. In our study, more VIP cells were activated after the extra similar and dissimilar conditions, compared to the control condition in which none of the objects were moved. In our study, it appears that MEC VIP cells may receive a novelty signal, enabling the activation of different populations of excitatory cells within the superficial layers of the MEC. Specifically, layer II stellate cells, which form synapses with both DG and CA3 cells^23,53^ and are essential for recognizing novel object location^23^, emerge as potential targets for VIP-mediated disinhibition. Hypothetically, if VIP interneurons participate in the disinhibition of grid cells, one of the functions in part supported by stellate cells^54,55^, this could lead to a greater strength of these neurons firing into downstream hippocampal cells, increasing the excitability of these neurons. Moreover, a portion of stellate cells have been shown to express head-direction activity^55^, known to exert a strong influence on grid cell activity^10,11^. However, the specific connectivity pattern between VIP cells and stellate cells remains to be determined.

More detailed investigations are needed to reconcile the plausible role of VIP cells in the modulation of cortical head-direction signals and their implication in spatial memory. Determining (1) if and through which connectivity scheme cortical VIP cells control head-direction cells activity and (2) how inhibition of cortical VIP cells interfere with memory in the MEC might participate in elucidating the functional role of this cell population.

### Technical limitations

Several technical limitations must be considered, as they might influence the results reported in this study. First, although many entorhinal cortical cell types were studied, our study covered select cellular subtypes. In particular, excitatory cells from MEC deep layers were not investigated here. Therefore, it is essential to consider that some studied regions potentially send most of their inputs to MEC deep layers, as it is already known for the retrosplenial cortex^56^. Second, using rabies-mediated retrograde labeling unilaterally, we found very few input cells in the contralateral hemisphere. Thus some anatomical pathways might have been overlooked, such as those involving stellate and pyramidal cells, which are typically known to receive substantial projections from the contralateral hemisphere^56,57^.

## Conclusion

This study corroborates similar results obtained in various brain regions supporting the idea that VIP cells are important to relay information provided by long-range inputs into the local network. More specifically, this study reveals the existence of a direct thalamocortical pathway to the MEC through VIP cells that could support essential computations in the head-direction system. Importantly, we demonstrate that this cell population is involved in spatial memory and could be essential for detecting environmental changes. Altogether, this work sheds light on the traditionally understudied MEC VIP cells and provides new insights into their contribution to MEC function, spatial cognition, and memory.

## Materials and Methods

### Animals

This study was carried out following the recommendations of the Canadian Council on Animal Care and the McGill University Animal Care Committee. The McGill University Animal Care Committee approved the protocol. Animals were housed in a temperature-controlled room with a 12/12 h light/dark cycle and food and water *ad libitum*. Homozygous Vip-IRES-cre (VIP^cre^, #010908, The Jackson Laboratory) mice were crossed with homozygous Ai9 lox-stop-lox-tdTomato cre-reporter mice (#007905, The Jackson Laboratory) to generate VIP-tdTom mice in which tdTom (Tom) is exclusively expressed in cells that have cre recombinase. Dr. Matthew Nolan, University of Edinburgh, kindly gave sim1^cre^ mice (C57Bl6/J). Sst-IRES-cre (SOM^cre^, #018973), Pvalb-IRES-cre (PV^cre^, #017320), and pOxr1-cre (pOx^cre^, #030484) were purchased from The Jackson Laboratory.

### Stereotaxic surgeries for viral transfection

5-8 weeks-old mice were anesthetized, their body temperature was maintained with a heating pad, and their eyes were hydrated with gel (Optixcare). Carprofen (2 mg/g) and saline (0.5 ml) were administered subcutaneously during the surgery. For each craniotomy, two injection sites were used. For the first site, the injection pipette was lowered 3.65mm from the bregma,; for the 2^nd^ site, it was lowered to 3.45mm from the bregma. The pipette was moved 0.280mm anterior to the transverse sinus for both injection sites to target the MEC. The injection depth was determined by the extent to which the micropipette bent once it hit the dura lining surface of the brain. Approximate depths for 5-week-old mice were: 1^st^ injection: −2.240, 2^nd^ injection site: −3.00. For each injection site, the pipette was retracted 200µm after the first bend, and the virus was injected at three different depths along the dorsal-ventral at 300µm intervals along the extent of the MEC, such that the last injection site usually corresponded to −1.450 to −1.750 along the DV axis. After each injection, the micropipette was left undisturbed for 3-4 minutes to allow for the infusion of the virus before injecting into the next site. A total of 400µl of virus was injected into each animal.

### Injection timeline for rabies-mediated viral tracing

VIP^cre^ (n = 4), sim1^cre^ (n = 3), pOx^cre^ (n = 4), PV^cre^ (n = 2), and SOM^cre^ (n = 4) mice were used for tracing monosynaptic inputs to MEC. These animals were first unilaterally injected with a 1:1 combination of AAV2/8-hSyn-FLEX-TVA-P2A-eGFP-2A-oG (ULaval Vector Core). 4 weeks later, animals were anesthetized as previously described, and RABV-DeltaG-EnvA-mCherry virus was injected using scar on the skull as a guide for location. One week post-injection, animals were perfused (4% PFA) to detect inputs and starter cells.

### Injection timeline for chemogenetic viral expression

For chemogenetic experiments, (n = 15) VIP^cre^ mice were injected bilaterally or unilaterally with the cre-mediated Designer Receptor Exclusively Activated by Designer Drugs (DREADDs) viral construct AAV8.hSyn.DIO.hM4Di.mCherry (Addgene, 4.3e12 GC-ml, diluted to a third in sterile PBS) using the injection strategy as described above.

### Habituation and handling for behavior experiment

#### c-Fos experiments

One week before testing in the novel object location paradigm, VIPtdtom (n = 14) mice were handled daily for 3-5 minutes for five consecutive days.

#### Chemogenetic experiments

Animals were subjected to behavioral testing only after recovering from surgeries (2 weeks), followed by two weeks of handling. For the first week, animals were handled for 3-5 minutes. For the second week, they were restrained by the scruff, and the animals were administered i.p. injections with saline every couple of days.

#### Novel object location paradigm

To test spatial memory, we adapted a novel object location paradigm from van Goethem, van Hagen, and Prickaerts 2018. Briefly, mice were placed in an opaque circular open field measuring (40 cm in diameter) surrounded by black walls with salient high-contrast distal cues. During the four days of habituation, animals were placed daily 10 min in the open field. During the last two days of habituation, a set of 2 new and different objects was introduced in the open field. On day 5, animals were placed for 15 min in the open field (sampling) and reintroduced 90 min later for 6 min (test). During the sampling, two identical objects were aligned horizontally in the center of the open field. During the test, one object was moved to a novel location (dissimilar: 12 cm from the original location; extra-similar: 3 cm from the original location) while the other remained in its original location. The location and choice of the objects were counterbalanced between different groups. For both tasks, the time spent exploring each object in the sample and test phases was measured, and animals that did not explore at least 10 seconds for each object during the sampling phase were excluded from the analysis. In the control group, the two objects were left unmoved during the test phase. The discrimination ratio was calculated as follows: Exploration time_moved_– Exploration time_unmoved_)/(Exploration time_moved_+ Exploration time_unmoved_) (s). Two experimenters scored the exploration times manually, and the average of their exploration times was used for analysis.

#### c-Fos experiment

VIPtdTom animals were used and perfused (4% PFA) 90 mins after the test session, and their brains were serially sectioned for c-Fos staining.

#### Chemogenetic experiments

VIP^cre^ mice were injected with AAV8.hSyn.DIO.hM4Di.mCherry virus in the MEC. VIP^cre^ mice in the control group did not receive any viral injection. JHU37160 (JHU, HelloBio #HB6261, 1mg/kg, diluted in saline, ^49^) or saline was injected 30 mins before sampling. Each animal was tested twice, once with saline and once with JHU 2 weeks apart. Mice were then perfused (4%PFA) after testing, and the brains were sectioned to confirm viral expression and specificity of the injection location.

#### Immunohistochemistry

Mice were deeply anesthetized and intracardially perfused with 4% PFA in PBS. Fixed brains were snap-frozen in dry ice-chilled isopentane before being cut into 35-μm-thick sections using a cryostat (Leica Microsystems).

#### Rabies injections

For circuit mapping experiments, whole brains from injected animals were sliced in the horizontal and sagittal plane to retrieve whole brain sections. Post-slicing, the sections were rinsed in 1X PBS (3×5 min).

#### c-Fos immunostaining

The sections were rinsed in 1X PBS (3×10 min) and then incubated in 1X PBS containing 0.3% Triton-X with the c-Fos antibody (1:1000, Abcam AB222699) at room temperature for 24 hours. After one rinse in 1X PBS, the slices were incubated with the secondary antibody (1:1000, ThermoFisher Scientific #A21206, Alexa Fluor 488) for 2 hrs at RT and finally rinsed 3×10 min in 1X PBS. DAPI mounting medium was used as a counterstaining.

#### Tracing of input regions for rabies circuit tracing experiments

For visualization, analysis, and quantification purposes, the sections from the rabies experiment were imaged with a VS-120 Olympus slide scanner under epifluorescence in DAPI, FITC, and TRITC channels using an optical filter set from Semrock (Part Number: DA/FI/TR 4X4M-C-000), an Orca r2 Hamamatsu monochrome camera, and an Olympus ×10 objective (Molecular and Cellular Microscopy Platform—Douglas Research Center, Montreal, Canada). Virtual Slide Images (.vsi) files acquired from the slide scanner were opened in FIJI using the BIOP VSI Reader (EPFL, Lausanne, Switzerland) and exported as individual TIFF files for further analysis in FIJI (Image J version 2.1.0) and QuPath (Software version 0.4.3.1).

#### Mouse brain atlas and regional analysis

The correct localization of input cells (mCherry+) in the MEC was validated using ABBA (Aligning Big Brain Atlas - EPFL, Lausanne (https://biop.github.io/ijp-imagetoatlas/) to register and align the Allen Atlas using both affine and spline transformation on sagittal and horizontal sections. The output was imported from ABBA, and manual counting was performed in the region of interest (ROI) outlined using the Allen Atlas in QuPath. The convergence index was defined as the total number of inputs targeting a single starter cell (Total number of inputs for an ROI/Total number of starter cells). The fraction of inputs was defined as the percentage of inputs a brain region sends to starter cells in the MEC (Total number of inputs for an ROI/Total number of presynaptic inputs in the entire brain) for each animal.

#### Image acquisition and Data analysis for c-Fos quantification

VIP, c-Fos, and VIP/c-Fos+ cells were quantified using the optical fractionator method in StereoInvestigator software with a Zeiss ApoTome structured illumination device on a widefield microscope. Contours were drawn to delineate the MEC’s superficial (L1-III) and deep layers (LIV-VI) using DAPI as a reference. Using live counting, an experimenter blind to the experimental conditions counted all markers according to optical dissector inclusion-exclusion criteria at each cell’s widest point. Separate markers were used to count the number of Tom (VIP), c-Fos (GFP), and co-localized or double-labeled cells (yellow).

#### Data analysis and statistics

Statistical analyses were performed with GraphPad Prism 8. Unpaired Student t tests (Mann Whitney U), one-way analysis of variance (ANOVA), or two-way ANOVAs, with Tukey or Bonferroni multiple comparison corrections were used. Data are presented as mean ± SEM with individual data points. Differences were considered significant when p<0.05. Additionally, column statistics against a theoretical mean of 0 were performed on the behavioral analysis; if α=0.05, the animal was considered to have learned during the paradigm.

## Acknowledgments

We thank Dr. Justin Quinn Lee, Dr. Jennifer C.Robinson, Dr. Coralie-Anne Mosser, Etienne Maes, Harshith Nagaraj, Samantha L Rosa and Feikai Lin for comments on earlier versions of this manuscript. The present study used the services of the Molecular and Cellular Microscopy Platform in the Douglas Hospital Research Center.

## Author Contributions

MO and SB participated in designing and conducting all experiments, analyzing data, drafting and writing the manuscript; RSM participated in cell counting; MPB reviewed the manuscript and provided funding for the project.

---

**Supplementary Figure 1:**
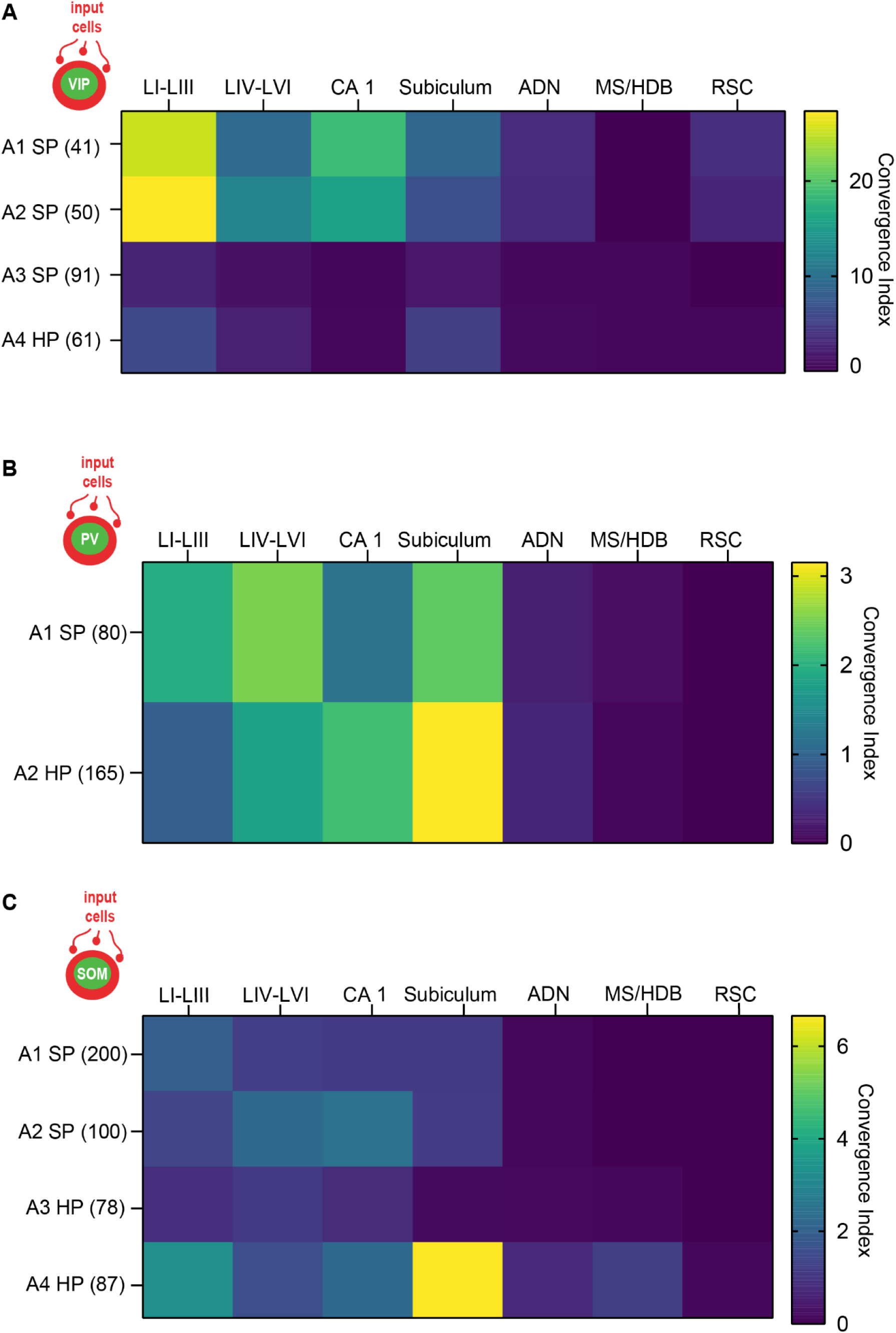
Quantification of inputs to VIP, PV, and SOM interneurons from individual mice. A. Input quantification for each VIP^cre^ mouse injected with rabies-mediated retrograde virus. B. Input quantification for each PV^cre^ mouse injected with rabies-mediated retrograde virus. C. Input quantification for each SOM^cre^ mouse injected with rabies-mediated retrograde virus. The number in the bracket indicates the number of starter cells quantified per individual animal. SP: sagittal plane; HP: horizontal plane.

**Supplementary Figure 2:**
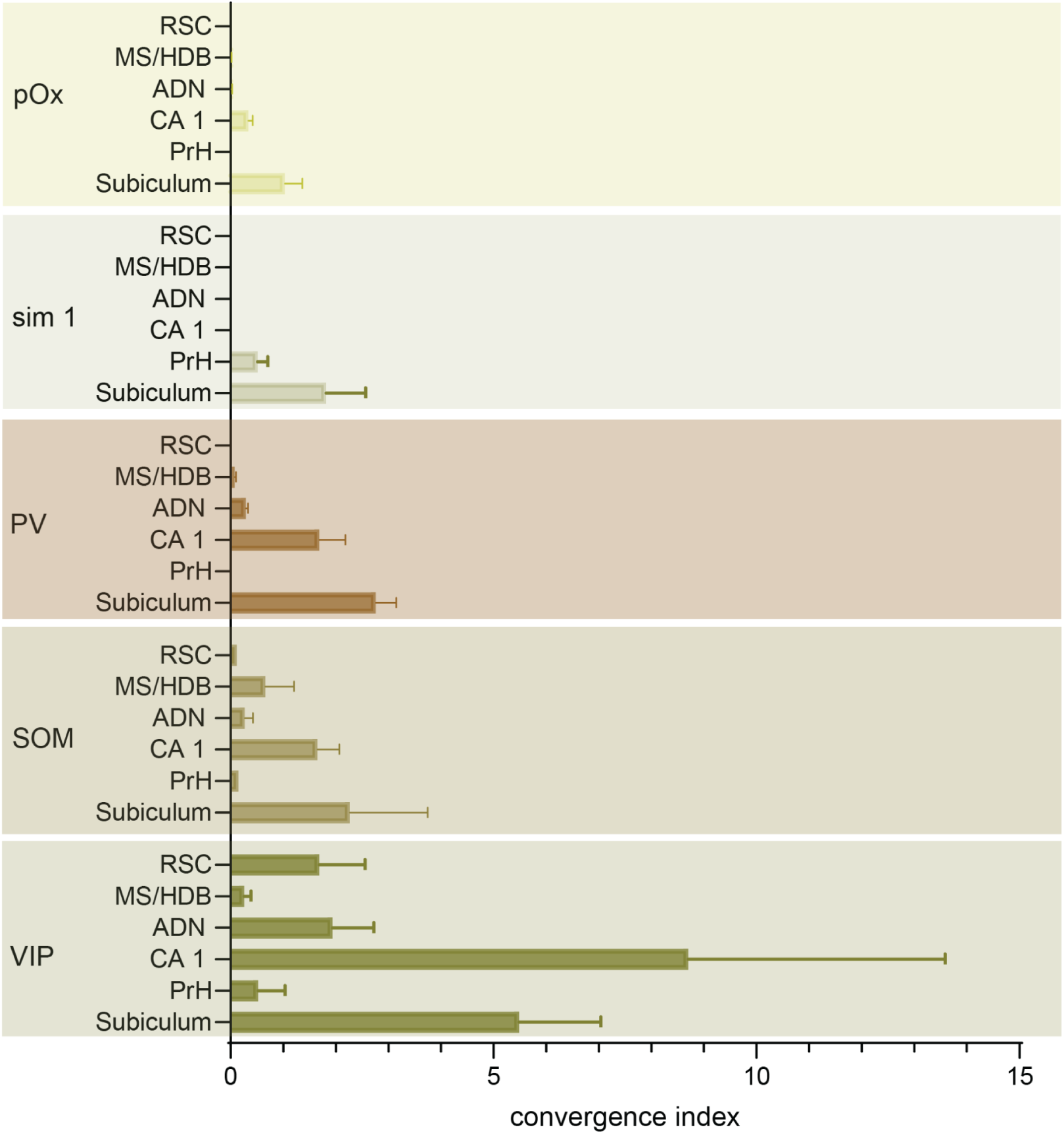
Comparative quantification of major inputs to all cell populations investigated RSC: retrosplenial cortex, MS/HDB: medial septum/horizontal diagonal band, ADN: anterodorsal nucleus of the thalamus, PrH: perirhinal cortex.

**Supplementary Figure 3:**
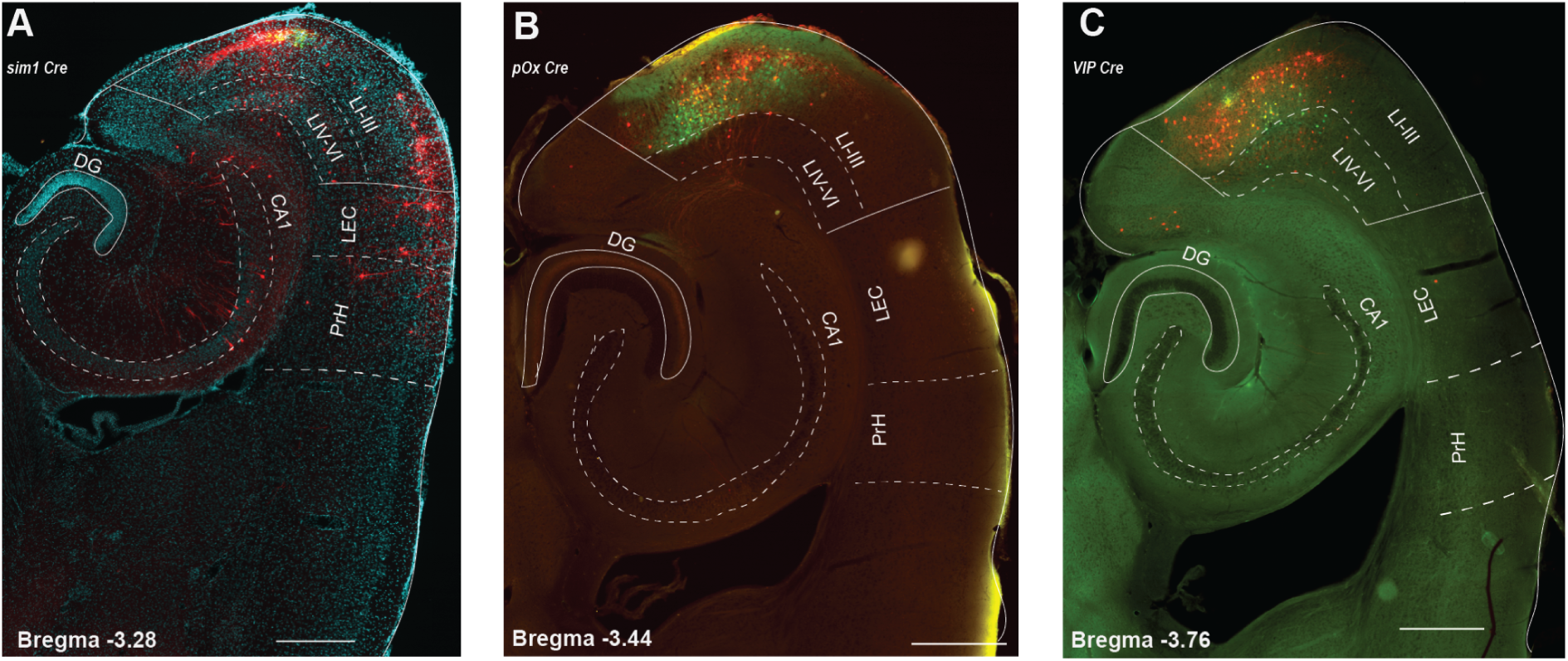
Input cells from the LEC are sparse compared to MEC cells Representative images of input cells in the LEC of sim1^cre^, pOx^cre^ animals and VIP^cre^ animals. In all images, mCherry are input cells, and GFP are TVA cells injected in the MEC. The scale bar is 500µm.

**Supplementary Figure 4:**
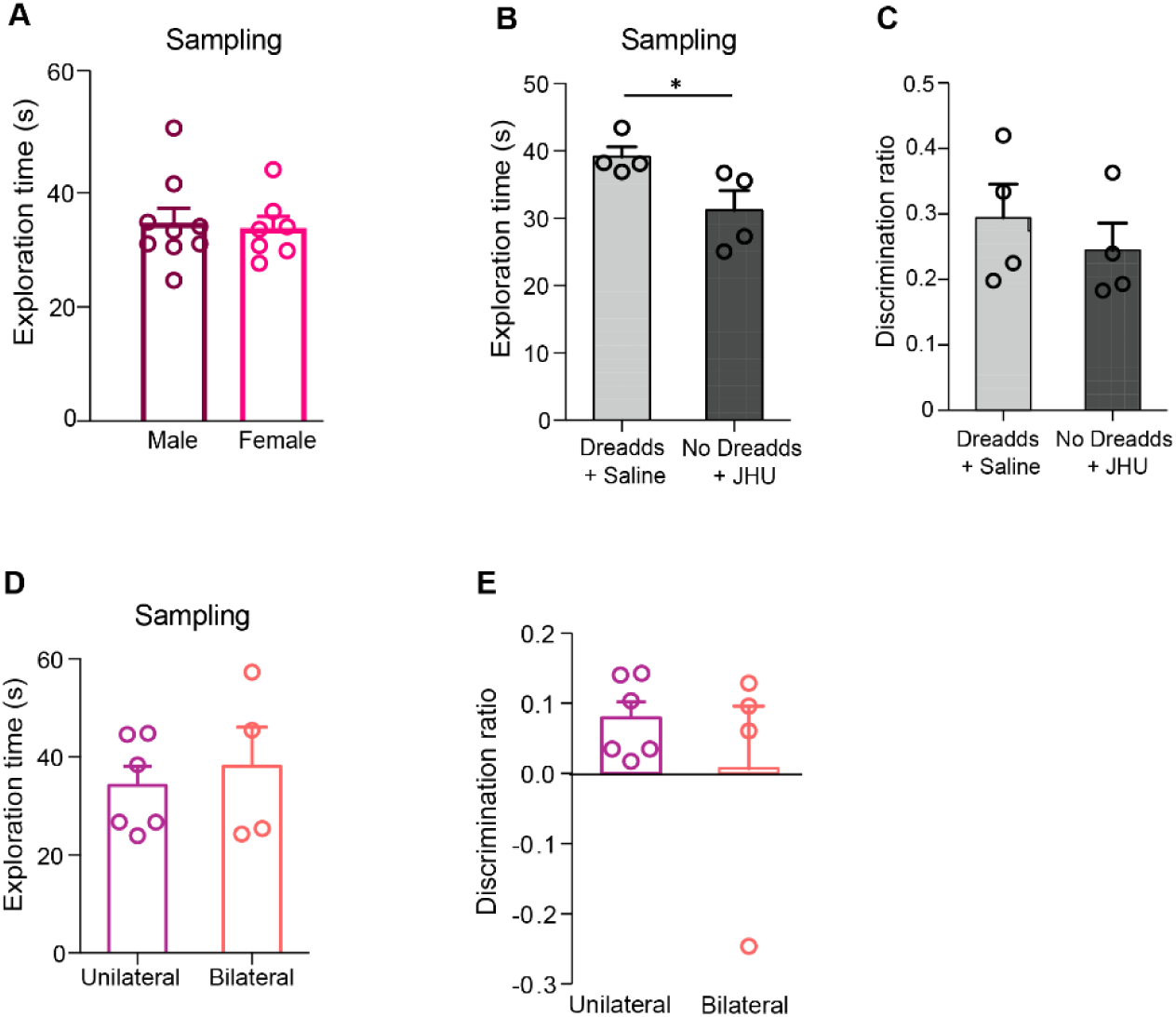
Quantification of the behavior in the NOL task A. Males (n = 9) and females (n = 7) spent equal time exploring the objects during the sampling phase of the NOL task. B. Mice injected with the AAV8-DIO-hSyn-hM4Di-mCherry virus that received saline (n = 4) spent more time exploring the object than mice with no viral injection but received JHU (n = 4) during the sampling phase of the NOL task (*p<0.05). C. No difference in the discrimination ratio of DREADDs + saline (n = 4) and no DREADDs + JHU (n = 4) was observed in the NOL task. D. No difference in the exploration time during the sampling phase of the NOL task was observed between animals that received unilateral (n = 6) or bilateral (n = 4) injections of the DREADD virus AAV8-DIO-hSyn-hM4Di-mCherry. E. No differences in the discrimination ratio were observed between animals that received unilateral (n = 6) or bilateral (n = 4) injections of the DREADD virus AAV8-DIO-hSyn-hM4Di-mCherry.

**Supplementary Table 1.** Tabulation of the number of cells counted in different regions for each cell line. In the table, PrH - Perirhinal cortex, ADN - Anterodorsal thalamic nuclei, MS/HDB - Medial Septum/Horizontal Diagonal Band of Broca and RSC - Retrosplenial Cortex.

| Mouse line | Animal ID | Plane of slicing | Starter cells |  | Projections |  |  |  |  |  |  |  |
| --- | --- | --- | --- | --- | --- | --- | --- | --- | --- | --- | --- | --- |
|  |  | SP - Sagittal<br>HP - Horizontal | Layer 1-3 | Layer 4-6 | MEC |  | Subiculum | PrH | Hippocampus | AD N | MS/HDB | RSC |
|  |  |  |  |  | Layers 1-3 | Layer 4-6 |  |  |  |  |  |  |
| VIP | Animal 1 | SP | 31 | 10 | 826 | 380 | 364 | 0 | 765 | 137 | 14 | 83 |
|  | Animal 2 | SP | 33 | 70 | 957 | 431 | 278 | 3 | 652 | 114 | 13 | 97 |
|  | Animal 3 | SP | 78 | 33 | 290 | 123 | 168 | 4 | 33 | 48 | 48 | 0 |
|  | Animal 4 | HP | 53 | 8 | 368 | 148 | 301 | 1 | 20 | 43 | 32 | 27 |
| PV | Animal 1 | HP | 48 | 32 | 157 | 75 | 190 | 0 | 95 | 20 | 8 | 0 |
|  | Animal 2 | HP | 89 | 76 | 417 | 299 | 520 | 0 | 361 | 55 | 8 | 0 |
| SOM | Animal 1 | SP | 157 | 43 | 360 | 239 | 218 | 0 | 216 | 22 | 0 | 10 |
|  | Animal 2 | SP | 39 | 61 | 138 | 224 | 114 | 0 | 250 | 8 | 0 | 0 |
|  | Animal 3 | SP | 35 | 43 | 68 | 85 | 14 | 0 | 66 | 11 | 9 | 10 |
|  | Animal 4 | HP | 46 | 41 | 287 | 137 | 579 | 0 | 193 | 64 | 105 | 1 |
| pOx | Animal 1 | SP | 82 | 10 | 42 | 18 | 130 | 0 | 41 | 4 | 0 | 0 |
|  | Animal 2 | SP | 155 | 17 | 346 | 208 | 141 | 0 | 86 | 2 | 0 | 0 |
|  | Animal 3 | HP | 50 | 0 | 225 | 0 | 85 | 0 | 6 | 1 | 2 | 0 |
|  | Animal 4 | HP | 127 | 0 | 99 | 24 | 20 | 0 | 30 | 1 | 1 | 0 |
| sim 1 | Animal 1 | SP | 27 | 0 | 121 | 28 | 23 | 19 | 1 | 0 | 0 | 0 |
|  | Animal 2 | SP | 47 | 0 | 249 | 162 | 61 | 15 | 0 | 0 | 0 | 0 |
|  | Animal 3 | HP | 112 | 0 | 177 | 71 | 369 | 0 | 0 | 0 | 0 | 0 |

## References

1. Tukker, J. J. et al. Microcircuits for spatial coding in the medial entorhinal cortex. Physiol. Rev. 102, 653–688 (2022).

2. Moser, E. I., Kropff, E. & Moser, M.-B. Place cells, grid cells, and the brain’s spatial representation system. Annu. Rev. Neurosci. 31, 69–89 (2008).

3. Sasaki, T., Leutgeb, S. & Leutgeb, J. K. Spatial and memory circuits in the medial entorhinal cortex. Curr. Opin. Neurobiol. 32, 16–23 (2015).

4. Hafting, T., Fyhn, M., Molden, S., Moser, M.-B. & Moser, E. I. Microstructure of a spatial map in the entorhinal cortex. Nature 436, 801–806 (2005).

5. Savelli, F., Yoganarasimha, D. & Knierim, J. J. Influence of boundary removal on the spatial representations of the medial entorhinal cortex. Hippocampus 18, 1270–1282 (2008).

6. Evans, T., Bicanski, A., Bush, D. & Burgess, N. How environment and self-motion combine in neural representations of space. J. Physiol. 594, 6535–6546 (2016).

7. Giocomo, L. M. et al. Topography of head direction cells in medial entorhinal cortex. Curr. Biol. 24, 252–262 (2014).

8. Sargolini, F. et al. Conjunctive representation of position, direction, and velocity in entorhinal cortex. Science 312, 758–762 (2006).

9. Høydal, Ø. A., Skytøen, E. R., Andersson, S. O., Moser, M.-B. & Moser, E. I. Object-vector coding in the medial entorhinal cortex. Nature 568, 400–404 (2019).

10. Gerlei, K. et al. Grid cells are modulated by local head direction. Nat. Commun. 11, 4228 (2020).

11. Grieves, R. M. & Jeffery, K. J. The representation of space in the brain. Behav. Processes 135, 113–131 (2017).

12. Ajabi, Z., Keinath, A. T., Wei, X.-X. & Brandon, M. P. Population dynamics of head-direction neurons during drift and reorientation. Nature 615, 892–899 (2023).

13. Taube, J. S. Head direction cells recorded in the anterior thalamic nuclei of freely moving rats. J. Neurosci. 15, 70–86 (1995).

14. Jacob, P.-Y. et al. An independent, landmark-dominated head-direction signal in dysgranular retrosplenial cortex. Nat. Neurosci. 20, 173–175 (2017).

15. Simonnet, J. & Fricker, D. Cellular components and circuitry of the presubiculum and its functional role in the head direction system. Cell Tissue Res. 373, 541–556 (2018).

16. Simonnet, J. et al. Activity dependent feedback inhibition may maintain head direction signals in mouse presubiculum. Nat. Commun. 8, 16032 (2017).

17. Clark, B. J. & Taube, J. S. Vestibular and attractor network basis of the head direction cell signal in subcortical circuits. Front. Neural Circuits 6, 7 (2012).

18. Clark, B. J. et al. Comparison of head direction cell firing characteristics across thalamo-parahippocampal circuitry. Hippocampus 34, 168–196 (2024).

19. Taube, J. S. The head direction signal: origins and sensory-motor integration. Annu. Rev. Neurosci. 30, 181–207 (2007).

20. Golob, E. J., Wolk, D. A. & Taube, J. S. Recordings of postsubiculum head direction cells following lesions of the laterodorsal thalamic nucleus. Brain Res. 780, 9–19 (1998).

21. Preston-Ferrer, P., Coletta, S., Frey, M. & Burgalossi, A. Anatomical organization of presubicular head-direction circuits. Elife 5, (2016).

22. Van Groen, T. & Wyss, J. M. Projections from the anterodorsal and anteroventral nucleus of the thalamus to the limbic cortex in the rat. J. Comp. Neurol. 358, 584–604 (1995).

23. Tennant, S. A. et al. Stellate Cells in the Medial Entorhinal Cortex Are Required for Spatial Learning. Cell Rep. 22, 1313–1324 (2018).

24. Vandrey, B. et al. Fan Cells in Layer 2 of the Lateral Entorhinal Cortex Are Critical for Episodic-like Memory. Curr. Biol. 30, 169–175.e5 (2020).

25. Kitamura, T. et al. Entorhinal Cortical Ocean Cells Encode Specific Contexts and Drive Context-Specific Fear Memory. Neuron 87, 1317–1331 (2015).

26. Alonso, A. & Klink, R. Differential electroresponsiveness of stellate and pyramidal-like cells of medial entorhinal cortex layer II. J. Neurophysiol. 70, 128–143 (1993).

27. Beed, P. et al. Layer 3 Pyramidal Cells in the Medial Entorhinal Cortex Orchestrate Up-Down States and Entrain the Deep Layers Differentially. Cell Rep. 33, 108470 (2020).

28. Miao, C., Cao, Q., Moser, M.-B. & Moser, E. I. Parvalbumin and Somatostatin Interneurons Control Different Space-Coding Networks in the Medial Entorhinal Cortex. Cell 171, 507–521.e17 (2017).

29. Grosser, S. et al. Parvalbumin Interneurons Are Differentially Connected to Principal Cells in Inhibitory Feedback Microcircuits along the Dorsoventral Axis of the Medial Entorhinal Cortex. eNeuro 8, (2021).

30. Fernandez, F. R., Via, G., Canavier, C. C. & White, J. A. Kinetics and Connectivity Properties of Parvalbumin- and Somatostatin-Positive Inhibition in Layer 2/3 Medial Entorhinal Cortex. eNeuro 9, (2022).

31. Buetfering, C., Allen, K. & Monyer, H. Parvalbumin interneurons provide grid cell-driven recurrent inhibition in the medial entorhinal cortex. Nat. Neurosci. 17, 710–718 (2014).

32. de Filippo, R., et al. Somatostatin interneurons activated by 5-HT receptor suppress slow oscillations in medial entorhinal cortex. Elife 10, (2021).

33. Badrinarayanan, S., Manseau, F., Williams, S. & Brandon, M. P. A Characterization of the Electrophysiological and Morphological Properties of Vasoactive Intestinal Peptide (VIP) Interneurons in the Medial Entorhinal Cortex (MEC). Front. Neural Circuits 15, 653116 (2021).

34. Gurgenidze, S. et al. Cell-Type Specific Inhibition Controls the High-Frequency Oscillations in the Medial Entorhinal Cortex. Int. J. Mol. Sci. 23, (2022).

35. Krabbe, S. et al. Adaptive disinhibitory gating by VIP interneurons permits associative learning. Nat. Neurosci. 22, 1834–1843 (2019).

36. Dávid, C., Schleicher, A., Zuschratter, W. & Staiger, J. F. The innervation of parvalbumin-containing interneurons by VIP-immunopositive interneurons in the primary somatosensory cortex of the adult rat. Eur. J. Neurosci. 25, 2329–2340 (2007).

37. Pi, H.-J. et al. Cortical interneurons that specialize in disinhibitory control. Nature 503, 521–524 (2013).

38. Pfeffer, C. K., Xue, M., He, M., Huang, Z. J. & Scanziani, M. Inhibition of inhibition in visual cortex: the logic of connections between molecularly distinct interneurons. Nat. Neurosci. 16, 1068–1076 (2013).

39. Turi, G. F. et al. Vasoactive Intestinal Polypeptide-Expressing Interneurons in the Hippocampus Support Goal-Oriented Spatial Learning. Neuron 101, 1150–1165.e8 (2019).

40. Leroy, F. et al. Enkephalin release from VIP interneurons in the hippocampal CA2/3a region mediates heterosynaptic plasticity and social memory. Mol. Psychiatry 27, 2879–2900 (2022).

41. Lee, A. T. et al. VIP Interneurons Contribute to Avoidance Behavior by Regulating Information Flow across Hippocampal-Prefrontal Networks. Neuron 102, 1223–1234.e4 (2019).

42. Ramos-Prats, A. et al. VIP-expressing interneurons in the anterior insular cortex contribute to sensory processing to regulate adaptive behavior. Cell Rep. 39, 110893 (2022).

43. Miettinen, M., Koivisto, E., Riekkinen, P. & Miettinen, R. Coexistence of parvalbumin and GABA in nonpyramidal neurons of the rat entorhinal cortex. Brain Res. 706, 113–122 (1996).

44. Wouterlood, F. G., Härtig, W., Brückner, G. & Witter, M. P. Parvalbumin-immunoreactive neurons in the entorhinal cortex of the rat: localization, morphology, connectivity and ultrastructure. J. Neurocytol. 24, 135–153 (1995).

45. Ennaceur, A. & Delacour, J. A new one-trial test for neurobiological studies of memory in rats. 1: Behavioral data. Behav. Brain Res. 31, 47–59 (1988).

46. van Goethem, N. P., van Hagen, B. T. J. & Prickaerts, J. Assessing spatial pattern separation in rodents using the object pattern separation task. Nat. Protoc. 13, 1763–1792 (2018).

47. Save, E. & Sargolini, F. Disentangling the Role of the MEC and LEC in the Processing of Spatial and Non-Spatial Information: Contribution of Lesion Studies. Front. Syst. Neurosci. 11, 81 (2017).

48. Kinnavane, L., Amin, E., Olarte-Sánchez, C. M. & Aggleton, J. P. Medial temporal pathways for contextual learning: Network c-mapping in rats with or without perirhinal cortex lesions. Brain Neurosci Adv 1, (2017).

49. Bonaventura, J. et al. High-potency ligands for DREADD imaging and activation in rodents and monkeys. Nat. Commun. 10, 4627 (2019).

50. Nassar, M. et al. Anterior Thalamic Excitation and Feedforward Inhibition of Presubicular Neurons Projecting to Medial Entorhinal Cortex. J. Neurosci. 38, 6411–6425 (2018).

51. Guet-McCreight, A., Skinner, F. K. & Topolnik, L. Common Principles in Functional Organization of VIP/Calretinin Cell-Driven Disinhibitory Circuits Across Cortical Areas. Front. Neural Circuits 14, 32 (2020).

52. Allegra, M., Posani, L., Gómez-Ocádiz, R. & Schmidt-Hieber, C. Differential Relation between Neuronal and Behavioral Discrimination during Hippocampal Memory Encoding. Neuron 108, 1103–1112.e6 (2020).

53. Cholvin, T., Hainmueller, T. & Bartos, M. The hippocampus converts dynamic entorhinal inputs into stable spatial maps. Neuron 109, 3135–3148.e7 (2021).

54. Abstract. Preprint at 10.7554/elife.36664.001 (2018).

55. Rowland, D. C. et al. Functional properties of stellate cells in medial entorhinal cortex layer II. Elife 7, (2018).

56. Simonsen, Ø. W., Czajkowski, R. & Witter, M. P. Retrosplenial and subicular inputs converge on superficially projecting layer V neurons of medial entorhinal cortex. Brain Struct. Funct. 227, 2821–2837 (2022).

57. Tang, Q. et al. Anatomical Organization and Spatiotemporal Firing Patterns of Layer 3 Neurons in the Rat Medial Entorhinal Cortex. J. Neurosci. 35, 12346–12354 (2015).

